# TGFβ signaling is required for sclerotome resegmentation during development of the spinal column in *Gallus gallus*

**DOI:** 10.1101/2021.10.25.465780

**Authors:** Sade W. Clayton, Ronisha McCardell, Rosa Serra

## Abstract

We previously showed the importance of TGFβ signaling in development of the mouse axial skeleton. Here, we provide the first direct evidence that TGFβ signaling is required for resegmentation of the sclerotome using chick embryos. Lipophilic fluorescent tracers, DiO and DiD, were microinjected into adjacent somites of embryos treated with or without TGFβR1 inhibitor, SB431542, at developmental day E2.5 (HH16). Lineage tracing of labeled cells was observed over the course of 4 days until the completion of resegmentation at E6.5 (HH32). Vertebrae were malformed and intervertebral discs were small and misshapen in SB431542 injected embryos. Hypaxial myofibers were also increased in thickness after treatment with the inhibitor. Inhibition of TGFβ signaling resulted in alterations in resegmentation that ranged between full, partial, and slanted shifts in distribution of DiO or DiD labeled cells within vertebrae. Patterning of rostro- caudal markers within sclerotome was disrupted at E3.5 after treatment with SB431542 with rostral domains expressing both rostral and caudal markers. We propose that TGFβ signaling regulates rostro-caudal polarity and subsequent resegmentation in sclerotome during spinal column development.

## INTRODUCTION

Spinal column formation is a dynamic process that requires migration and subsequent differentiation of mesenchymal sclerotome cells into the vertebrae (VB), cartilaginous end plates, ribs, annulus fibrosus (AF) of the intervertebral discs (IVD), and tendons and ligaments of the spine (Christ et al., 2007, Alkhatib et al., 2018, Williams et al., 2019, Cox and Serra, 2014). This process begins with the formation of somites that will differentiate into dermomyotome and sclerotome, depending on the signals emanating from neighboring tissues (Kalcheim and Ben-Yair, 2005, Christ and Ordahl, 1995). These signals include Wnt1/3 from the epidermis and BMP4 from the lateral plate mesoderm causing differentiation of the dermomyotome from the lateral dorsal region of the somite while Shh and Noggin secreted from the notochord and floor plate of the neural tube stimulate sclerotome formation ventrally (Fan et al., 1997, Fan and Tessier-Lavigne, 1994, Marcelle et al., 1997).

Sclerotome is initially organized along the anterior-posterior axis of the embryo into a metameric pattern of rostro-caudal domains separated by von Ebner’s fissure (Christ et al., 2000, Von Ebner, 1888). The discrete rostral and caudal domains within each segment expresses distinct markers such as Tbx18, Mesp2, and Tenascin rostrally and Unxc4.1, Ripply 1/2, Pax 1/9, and Peanut Agglutinin (PNA) caudally (Christ and Ordahl, 1995, Kawamura et al., 2008, Neubuser et al., 1995, Leitges et al., 2000, Morimoto et al., 2007, Tan et al., 1987, Stern and Keynes, 1987). Development of the spinal column requires rostro-caudal polarization and then reorganization of the sclerotome, a process called resegmentation, to allow proper alignment of the spine with the tendon, musculature, and nerves (Huang et al., 2000, Williams et al., 2019, Cox and Serra, 2014, Alkhatib et al., 2018). The process of resegmentation is preceded by the formation of rostral and caudal domains within the sclerotome that regroup during resegmentation in response to stimuli that are still unknown (Remak, 1855, Bagnall et al., 1988, Huang et al., 2000). Rostral and caudal domains within each sclerotome segment separate and recombine with the corresponding adjacent segment. This results in the formation of a new sclerotome unit that is now shifted one half segment with respect to the myotome (Huang et al., 2000, Williams et al., 2019, Alkhatib et al., 2018, Cox and Serra, 2014). The newly resegmented sclerotome will develop into the VB, and the AF will develop in chick from cells in the rostral domain immediately adjacent to the original rostral-caudal border at von Ebner’s fissure (Bruggeman et al., 2012). Tendon will form from the cells adjacent to the myotome (Brent et al., 2003). The signaling pathways that regulate resegmentation are unknown.

TGFβ is a multifunctional growth factor that controls many aspects of development. TGFβ signals as a dimer that binds to a heterotetrametric receptor complex on the cell membrane that consists of two TGFβ type 1 receptors, TGFβR1, and two TGFβ type two receptors, TGFβR2. Activation of the receptor complex stimulates the phosphorylation of the well characterized downstream effectors Smad 2/3, or various “non-canonical” downstream effectors including ERK 1/2, AKT, and p38 (Hata and Chen, 2016, Chen et al., 2019, Zhang, 2009, Clayton et al., 2020). TGFβ regulates the expression of markers for fibrous tissues, including AF, ligament, and tendon, in cultured sclerotome through Smad-dependent and non-canonical signaling pathways (Clayton et al., 2020, Sohn et al., 2010, Ban et al., 2019, Cox et al., 2014). Deletion of *Tgfbr2* in mouse sclerotome *in vivo* (Col2aCre;*Tgfbr2*^LoxP/LoxP^) results in failure of the AF and other fibrous tissues to form correctly (Baffi et al., 2004, Pryce et al., 2009). In addition, loss of *Tgfbr2* results in phenotypes that would be consistent with defects in resegmentation including split lamina, disorganized costal joints, loss of the IVD, and defects in the anterior articular process ( Baffi et al., 2004, Baffi et al., 2006). Furthermore, *Tgbr2* deleted mice demonstrate alterations in the rostro-caudal polarity of the sclerotome with Pax1 and Pax9, markers of caudal sclerotome, being expressed through the entire segment (Baffi et al., 2006). Deletion of rostro-caudal markers Mesp2 and Rippy 1/2 in mice also contribute to alterations in the formation of the IVD (Takahashi et al., 2013). Alterations in rostro-caudal polarity and subsequent resegmentation would be expected to alter the context in which cells differentiate, affecting development of the spinal column.

Here, we provide the first direct evidence that TGFβ regulates resegmentation. By using a drug inhibitor, SB431542, to inhibit TGFβ signaling within the thoracic somites and lipophilic dyes, DiD and DiO, to lineage trace labeled cells from the somite, we show that TGFβ signaling is required for resegmentation of sclerotome. In addition, inhibition of TGFβ signaling resulted in mishappen VBs, reduced IVD, and alterations in rostro-caudal polarity of the sclerotome. We conclude that TGFβ regulates resegmentation and development of the spinal column.

## RESULTS

### SB431542 injection into chick somites disrupts TGFβ signaling in somite derivatives

Previous studies from our lab showed that TGFβR2 is required for the development of the mouse axial skeleton (Col2aCre;*Tgfbr2*^LoxP/LoxP^ mice; Baffi et al., 2004, Baffi et al., 2006). Some of the defects observed in the Col2aCre;*Tgfbr2^LoxP/LoxP^* mice could be consistent with defects in resegmentation (Baffi et al., 2006); however, this has not been directly tested. To directly test the role of TGFβ in resegmentation, we utilized the chick model since spinal column development is easily observed *in vivo.* First, we demonstrated the efficacy of a TGFβR1 inhibitor, SB431542, in chick embryos through determination of the expression pattern and level of Scx, a known downstream target of TGFβ signaling (Clayton et al., 2020). Embryonic day 2.5 (E2.5) chick embryos were injected with DMSO or SB431542 in paired thoracic somites 19-26 (Fig. 1A). One day later, E3.5, embryos injected with SB431542 showed a reduction in Scx mRNA by in situ hybridization specifically within sclerotome derived from the inhibitor injected somites (Fig. 1C, white arrows). Sclerotome derived from surrounding somites continued to express Scx mRNA (Fig. 1C) as did sclerotome derived from somites injected with the DMSO control (Fig. 1B). Next, tissue from the injected area was dissected from E3.5 embryos. Western blot analysis of protein lysates indicated a statistically significant reduction in Scx protein in SB431542 injected embryos when compared to DMSO controls (Fig. 1D, E). Protein levels were normalized to the loading control alpha tubulin (Fig. 1D, E). In addition, pSmad3, a direct effector of TGFβ signaling, and Adamtsl2, another downstream target of TGFβ, were downregulated in SB431542 treated somites relative to DMSO controls (Figure 1D). Reduced levels of TGFβ responsive targets in SB431542 treated somites indicated that the inhibitor was working *in vivo* at the concentrations used.

**Fig. 1:**
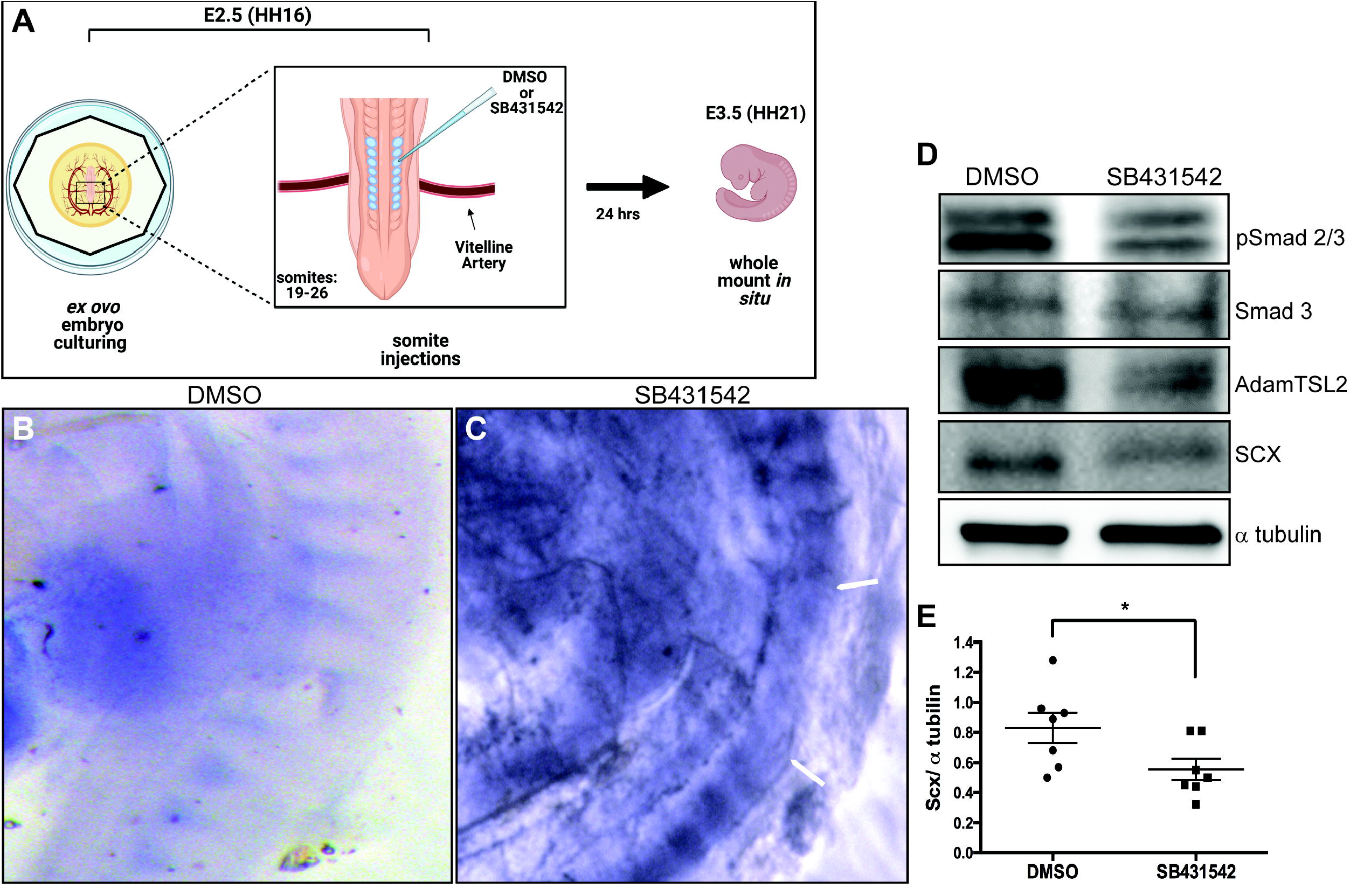
SB431542 treatment disrupts TGFβ signaling and formation of the spinal column in chick embryos. (A) Schematic of *ex ovo* culture and injection in chick embryos. (B, C) Whole mount *in situ* hybridization in E3.5 embryos with an antisense probe to Scx mRNA. (B) DMSO injected control embryos (n=5) or (C) SB431542 injected embryos (n=3). The white arrows in panel C denote the range of SB431542 injection. (D) Thoracic tissue was isolated from control and treated embryos and western blot analysis was conducted using pSmad3 (n=3), Smad3 (n=3), Scx (n=7), Adamtsl2 (n=2), and α tubulin (n=7) specific antibodies. α tubulin was used as a loading control. (E) Densitometry was used to quantify relative expression of Scx/ α tubulin from western blots (n=7). Differences between groups was analyzed using an unpaired two-tailed t-test in GraphPad Prism. n= denotes the number of biological replicates. * = p < 0.05. Left is ventral and right is dorsal, top is anterior and bottom is posterior in B and C.

We then wanted to determine if the chick model recapitulated the spinal defects seen in mice. Chicks that were injected with DMSO or SB431542 at E2.5 were harvested at E6.5 and E12.5 and stained with Alcian blue to highlight cartilage and skeletal development (Fig. 2). E6.5 chick embryos were sectioned, and midline sections were stained with Alcian blue (Fig 2A-D). Alcian blue stains sulfated glycosaminoglycans and glycoproteins and is a histological marker for cartilage (Nagy et al., 2009). Compared to control, the spinal column in SB431542 treated embryos demonstrated multiple defects (Fig 2. A-D). Vertebrae walls, red bars, were thinner (Fig. 2C), IVD disc height, black brackets, was reduced (Fig. 2D), and the rib heads, black arrows, were malformed in the SB431542 treated group compared to DMSO treated controls.

**Fig. 2:**
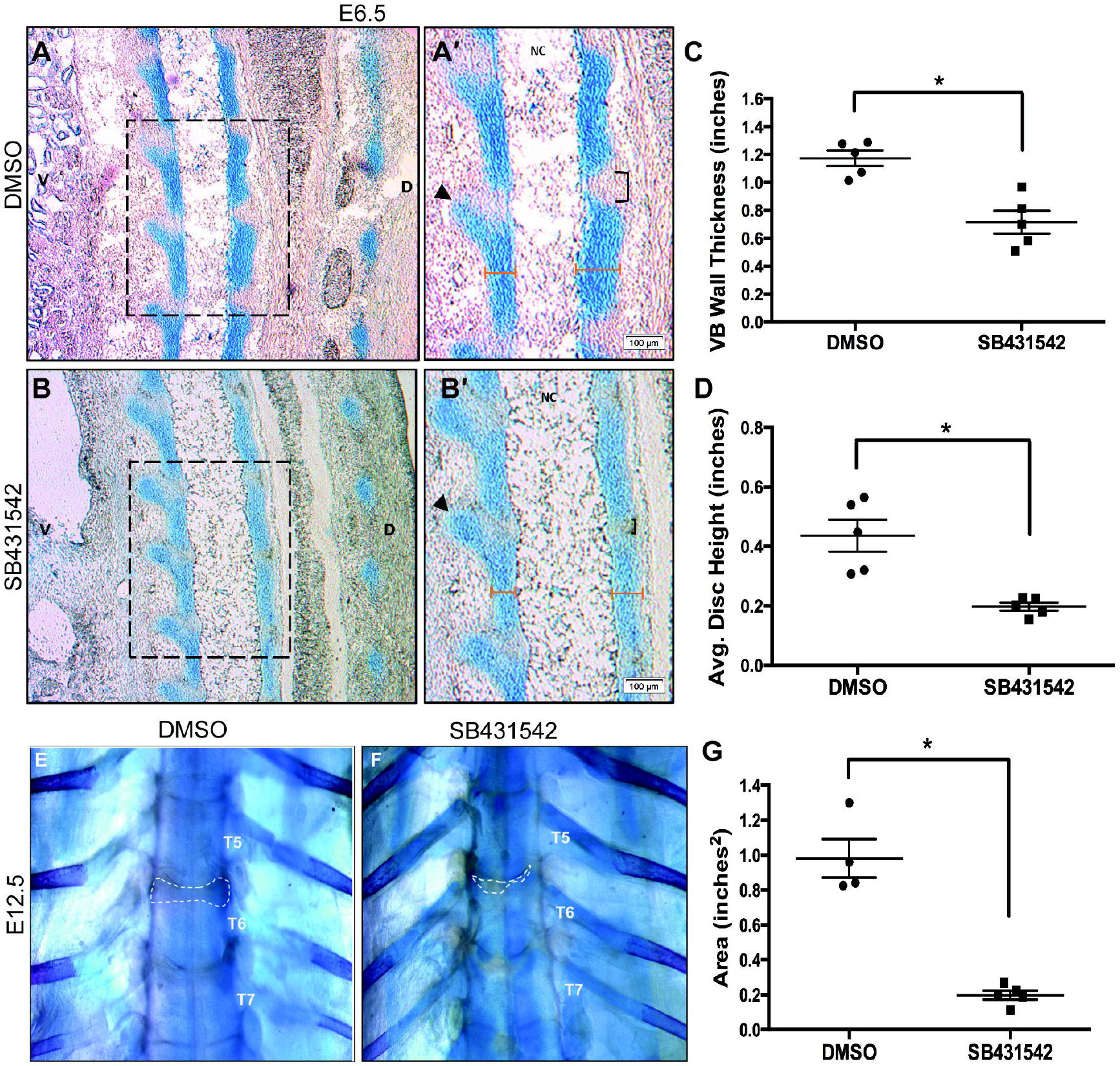
Spinal column development is altered when TGFβ signaling is inhibited. (A-D) E6.5 embryos were embedded in OCT, sectioned, and the midline sections that showed the notochord were stained with alcian blue. (A) Control embryos (n=5) or those treated with (B) SB431541 (n=5) showed (C) reduced vertebral wall thickness and (D) reduced IVD height. A’ and B’ are magnified regions from panels A and B. The black arrows in panels A’ and B’ point to the rib head. Black brackets indicate the disc space, and the red bars indicate the VB wall thickness. (EG) Skeletal preparations using alcian blue and alizarin red staining were performed on E12.5 embryos. The thoracic segment of (E) DMSO control (n=4) and (F) SB431542 treated (n=5) embryos were observed and the (G) IVD area was quantified. n= denotes the number of biological replicates. * = p < 0.05. NC, notochord; D, dorsal; V, ventral.

Next, skeletal preparations using Alcian blue and Alizarin red staining were performed on E12.5 embryos to analyze mature skeletons after inhibiting TGFβ signaling in somites. Control embryos had distinct oval shapes of the AF within the IVD space while inhibitor treated embryos displayed under formed and misshapen AF (Fig. 2E-G, outlined in white). Quantification of the average area of the AF (outlined in white) indicated that the AF in SB431542 injected embryos was reduced compared to controls (Fig. 2G), similar to what is observed in Col2aCre;*Tgfbr2*^LoxP/LoxP^ mice. The results indicated that TGFβ signaling is required for normal skeletal development in chick and validates the model for subsequent studies.

TGFβ signaling also been shown to act as an inhibitor of myofiber differentiation (Kollias and McDermott, 2008, Liu et al., 2001, Filvaroff et al., 1994). To further confirm the activity of SB431542 in chick embryos, we compared sections from control and SB431542 treated embryos for MF20 expression by immunofluorescent staining (Fig 3. A, B). The area between the ribs containing hypaxial muscle fibers in E6.5 embryos was compared. Fibers appeared thicker in the SB431542 treated embryos. This was confirmed through morphometric analysis (Fig. 3 C). There was a statistically significant increase in myofiber thickness in the SB431542 treated embryos. We then observed myocyte formation in real-time in labeled control and SB431542 treated embryos (Fig. S1, Movies 1, 2 and 3). The elongated myocytes appeared increased in number and thickness when compared to control embryos. Inhibition of TGFβ signaling caused increased myofiber differentiation in SB431542 injected embryos (Movie 3) relative to controls (Movie 2) further supporting the use of the model (Liu et al., 2001, Kollias and McDermott, 2008).

**Fig. 3:**
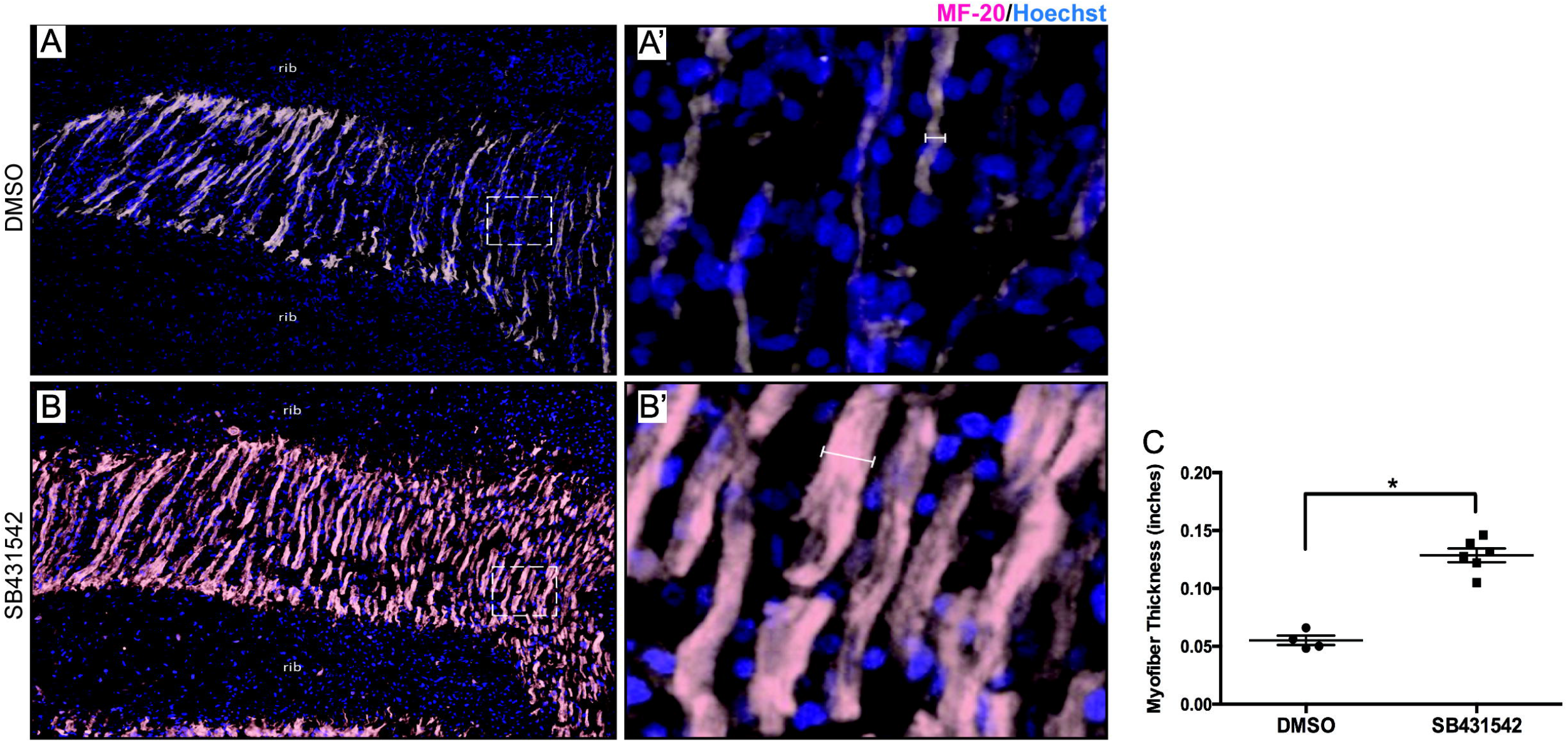
Myogenic differentiation is altered when TGFβ signaling is inhibited. (A,B) E6.5 embryos were embedded in OCT, sectioned, and slides containing ribs and hypaxial muscle fibers were selected and stained with MF20 (pink) and counter stained with Hoechst (blue). (B) DMSO injected controls (B’) zoomed in region from white box in B (n=4). (C) SB431542 injected embryos (C’) a zoomed in region from white box in C (n=6). (C) Average myofiber thickness (white brackets) was measured using ImageJ, and averages were analyzed using an unpaired two-tailed t-test in GraphPad Prism. * = p < 0.05. n= denotes the number of biological replicates. Left is ventral, and right is dorsal.

### TGFβ signaling is required for resegmentation

Many details of resegmentation in the sclerotome are still unknown (Christ et al., 2000, Goldstein and Kalcheim, 1992, Huang et al., 2000). To determine the time course of sclerotome resegmentation, we utilized lipophilic, fluorescent tracker dyes. DiD, far red, and DiO, green. These dyes are dialkylcarbocyanines that intercalate into the cell membrane and thus can be used as cell lineage tracers of any cells they come into contact with (Honig and Hume, 1989). At E2.5, thoracic somites 21 through 26, right side only, were injected in an alternating pattern of DiD, somites 21, 23 and 25, or DiO, somites 22, 24, and 26 (Fig. 4A). This alternating pattern was done to monitor resegmentation and show how somites eventually contribute to a particular vertebra (Ward et al., 2017). We chose the thoracic region somites 21-26 because: 1) the large vitelline artery crosses underneath somite 23 at this developmental stage and therefore, could be used as a reliable morphological marker to inject the same somites in every embryo, 2) these are newly formed somites at the time of injection and therefore should not have undergone differentiation, 3) since this is restricted to the thoracic region, they should have the same resegmentation pattern (Ward et al., 2017), and 4) vertebral fusion does not normally occur in the thoracic region of *Gallus gallus* until several weeks after hatching. Furthermore, the right axis of the embryo was injected since after the embryo turns the right side is facing up making imaging possible.

**Fig. 4:**
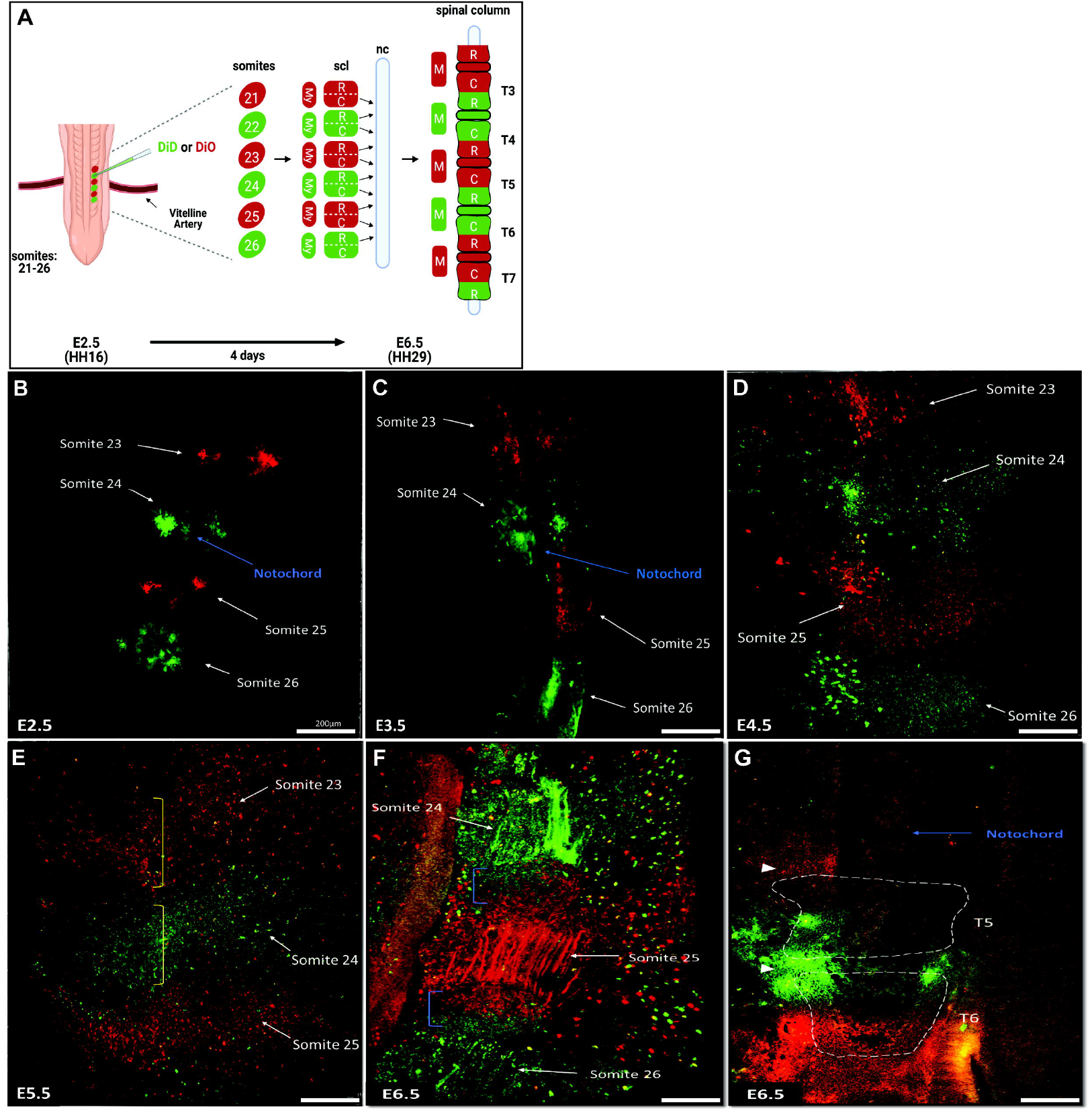
Resegmentation is completed in the chick spinal column by E6.5 days. (A) Schematic of lipophilic dye injections and the process of resegmentation in thoracic sclerotome. (B-F) Embryos were isolated after dye injections and 3D volumetric projections (maxIP images) of the embryos were acquired starting at (B) E2.5 days (n=3) and taken every 24hrs at (C) E3.5 (n=3), (D) E4.5 (n=3), (E) E5.5 (n=11) until (F) E6.5 (n= 6) where resegmentation was evident in the cartilaginous rib heads (F, blue brackets). (G) A section of an E6.5 embryo at the midline showed evidence of resegmentation in the VBs (outlined by white boxes) (n=5). The yellow brackets in panel E show where the labeled cells that have migrated ventrally have begun to segment. The white arrowheads in panel G show the IVD regions. n= denotes the number of biological replicates. Scl, sclerotome; NC, notochord; My, myotome; M, muscle; T, thoracic vertebrae; R, rostral; C, caudal. Dorsal is right. Ventral is left. Anterior is to the top and posterior to the bottom.

We first used max intensity projection (maxIP) 3D images to visualize DiD and DiO labeled cells within the spinal column. Max IP projections represented a single time point image that captures a volumetric snapshot through 150 to 200 microns of the sample. For imaging, embryos were injected with dyes at E2.5, isolated at the indicated times, embedded in a yolk/agarose imaging solution to maintain viability, and then imaged on a laser scanning confocal microscope. Excessive cell death as determined by Calcein-Ethidium cell staining was not observed in isolated embryos embedded in the yolk/agarose solution even after 12 hrs when compared to embryos allowed to develop normally in *ex ovo* culture (Fig. S2). One Max IP image was obtained each day (E2.5 to E6.5) after injection. At E2.5 days, max IP images showed that the somites were well labeled (Fig. 4B). By E3.5, the spherical shape of the somite became less distinct suggesting epithelial to mesenchymal transition of cells in the somite (Fig 3B, C; Fig. S3). In addition, labeled cells on the ventral side of the notochord suggested migration and formation of sclerotome at E3.5 (Fig 4C; Fig. S3). Growth of the embryo was rapid between E3.5 and E4.5/ E5.5. There was an increase in the length of the dorsal-ventral axis relative to the anterior-posterior axis of each sclerotome segment suggesting continued migration of cells ventrally (Fig 4C-E). At 4.5 and E5.5, the segmented alternately labeled pattern of the sclerotome in the ventral portion of the embryo could be seen (Fig. 4D, E). In addition, as the embryo grew, it increased in thickness so that by E4.5 only very lateral aspects of the spine (150 to 200 microns deep) could be imaged (Fig. 4D). At E6.5, evidence of resegmentation was observed in the lateral cartilages between muscle fibers (Fig 4F, blue brackets = cartilage) where half of each cartilage element was stained far red and half green. To better image the midline, E6.5 embryos that had been injected with DiD and DiO were sectioned and the vertebrae and IVDs were identified by counterstaining with rhodamine conjugated peanut agglutinin (PNA) (Rashid et al., 2020). Sections were simultaneously imaged for all three fluorophores (rhodamine, DiD, and DiO) and the developing vertebrae were outlined by using the ImageJ threshold tool to decrease background noise in PNA stained images and then the ROI function was used to outline the VB (Fig. S4). Each vertebra was labeled half far red and half green, clear evidence of resegmentation (Fig. 4G). Previously studies have shown a sharp border between red and green domains with little to no cell mixing (Stern and Keynes, 1987). Our data supported these previous observations by showing distinctly labeled red and green cell domains except in the most superficial areas, for example, the developing skin. In summary, labeled cells were followed each day (Fig. 4) and we found that resegmentation was completed by E6.5 with each developing vertebrae consisting of a half far red and a half green domain (Fig.4 F, G).

Next, to determine the role of TGFβ signaling in resegmentation, SB431542, a TGFβR1 inhibitor, or DMSO, was mixed with DiD or DiO in a 1:1 ratio and injected into somites 21-26 in an alternating pattern as described above (Fig.4A; Fig. 5A). Sections from control and treated embryos were counterstained with rhodamine conjugated Peanut Agglutinin (PNA) to localize the developing VB as described above (Fig. 5B-E, outlined in white dotted box). When compared to embryos injected with DMSO, SB431542 injected embryos showed altered resegmentation patterns (summarized in Fig. 5A). In control embryos, VB consisted of half far red and half green domains as expected for normal resegmentation (Fig. 5B, F, G). The border between the labeled domains is marked with a yellow dotted line. Embryos injected with SB431542 demonstrated changes in the ratio of far red and green labeled domains within the VB that manifested as full shift, partial shift, slanted border, or a mixture of these. Full shift was visualized as only one color in the VB (Fig. 5C). A partial shift was defined as a change from 50-50 in the ratio of the far red and green domains in the VB (Fig. 5D). SB431542 injected embryos that demonstrated a partial shift had an expanded caudal domain (Fig. 5D, G, F). Slanted border was defined as an alteration in the ratio of red and green domains but the border between the two was not straight across the VB (Fig. 5E). Slanted borders also resulted in an overall expanded caudal domain (Fig. 5E, F, G). Some embryos demonstrated more than one alteration and had both a partial shift and slanted borders (Fig. 5D). Quantification of the volume of each domain relative to the total segment indicated that resegmentation was significantly affected by blocking TGFβ signaling with SB431542 indicating a role for TGFβ in resegmentation.

**Fig. 5:**
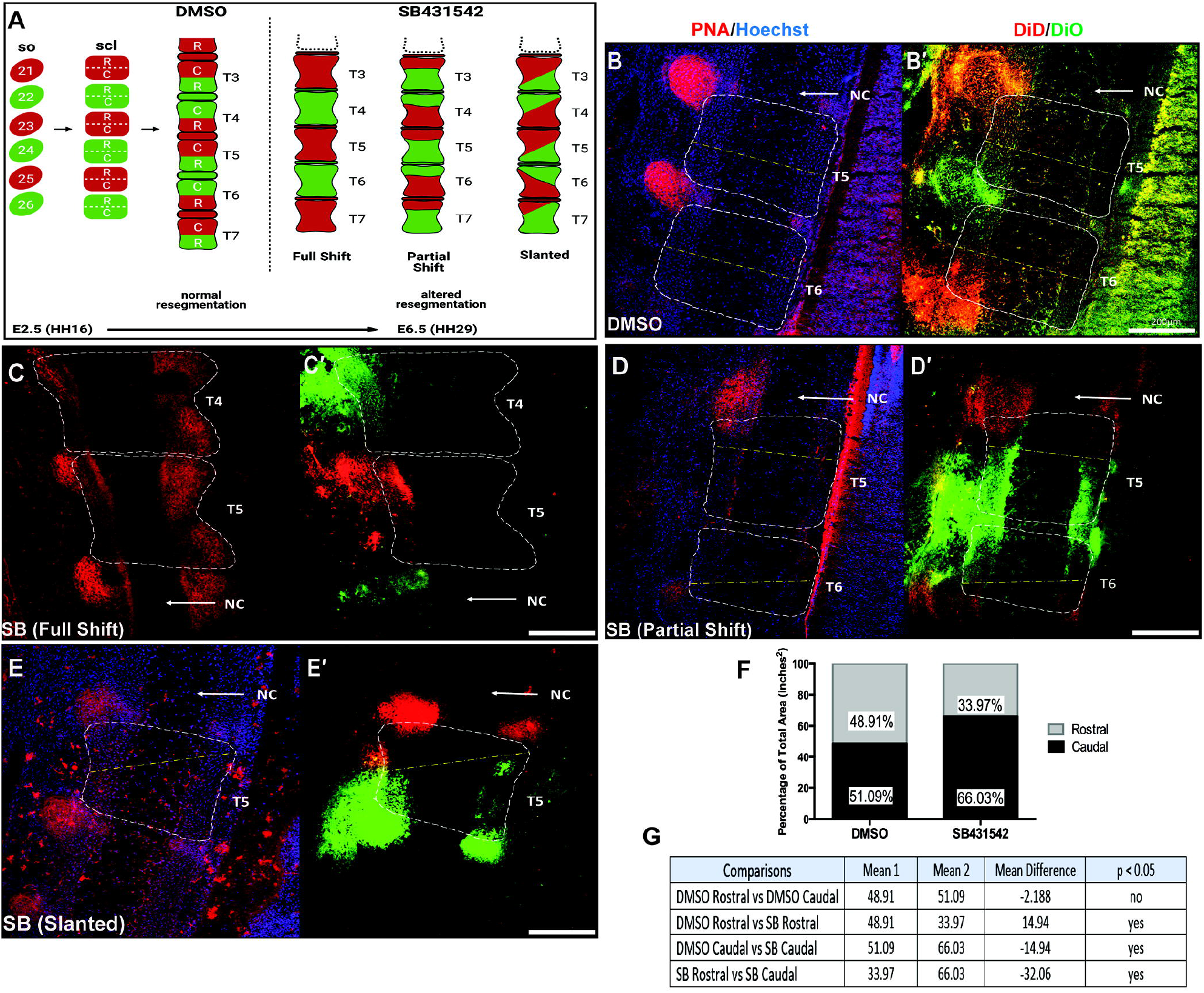
Inhibition of TGFβ signaling disrupts resegmentation. (A) Schematic of the process of normal resegmentation compared to altered resegmentation in SB431542 treated embryos at E6.5 days. SB431542 treated embryos were grouped into three different categories, full shift (n=3), partial shift (n=3), or slanted (n=3), based on the border between far red and green stained cells within the VB. (B) DMSO (n=5) or (C-E) SB431542 injected embryos were counterstained with peanut agglutin-RITC (PNA) and Hoechst and the vertebral body was outlined in white. The outlined VB area was then superimposed over the images of DiD/DiO labeled cells on the same slides (B’-E’). Examples of (C) a full shift (n=3), (D) partial shift (n=3), and (E) slanted shift (n=3) are shown. (F) An ANOVA analysis was used to compare the differences in the DiD/DiO labeled rostral and caudal zones in DMSO versus SB431542 vertebrae. (G) All comparisons are listed in the table. A Tukey’s post hoc analysis was used to compare differences between groups. VBs (white dotted squares) were outlined using the threshold and ROI tools on Image J. Rostro caudal boundary is denoted by a yellow dotted line. Left is ventral, and right is dorsal. n= denotes the number of biological replicates. So, somite; Scl, sclerotome; NC, notochord; T, thoracic vertebrae; R, rostral; C, caudal.

### TGFβ signaling regulates rostral-caudal patterning in the sclerotome

Sclerotome is organized into rostral and caudal domains that can be molecularly marked, for example, by Tenascin expression in the rostral domain and high PNA staining in the caudal domain (Huang et al., 2000, Stern and Keynes, 1987, Baffi et al., 2006, Cox and Serra, 2014, Alkhatib et al., 2018, Williams et al., 2019). The observations above suggested that SB431542 treated embryos had defects in resegmentation that resulted in an expansion of the caudal domain in each sclerotome segment. We previously noted that Col2aCre;*Tgfbr2*^LoxP/LoxP^ mice demonstrated expression of Pax1 and Pax9 through the entire sclerotome whereas Pax1/9 were only expressed in the caudal domain in control mice suggesting expansion of the caudal domain. Furthermore, rostralization or caudalization of sclerotome results in spinal phenotypes that are consistent with defects in resegmentation (Takahashi et al., 2013). To test the hypothesis that TGFβ signaling affects rostral-caudal polarity in the early chick sclerotome, embryos were injected with DMSO or SB431542 at E2.5 and markers for rostral (Tenascin) and caudal (PNA) domains of the sclerotome were localized in sections from E3.5 embryos using immunofluorescent staining (Fig.6). In DMSO treated embryos, Tenascin was localized to the rostral domain as expected and localization was comparable in SB431542 injected embryos (Fig6A, B). Next, sections were stained with PNA. In control embryos, staining was more intense in the caudal domain (Fig. 6C) as previously described (Stern and Keynes, 1987). In contrast, in SB431542 treated embryos PNA labeling was expanded throughout the rostral domain (Fig. 6D). These results along with our previous results in mouse (Baffi et al., 2006) suggest that TGFβ signaling regulates rostral-caudal polarity in sclerotome specifically by limiting the expansion of the caudal domain markers.

**Fig. 6:**
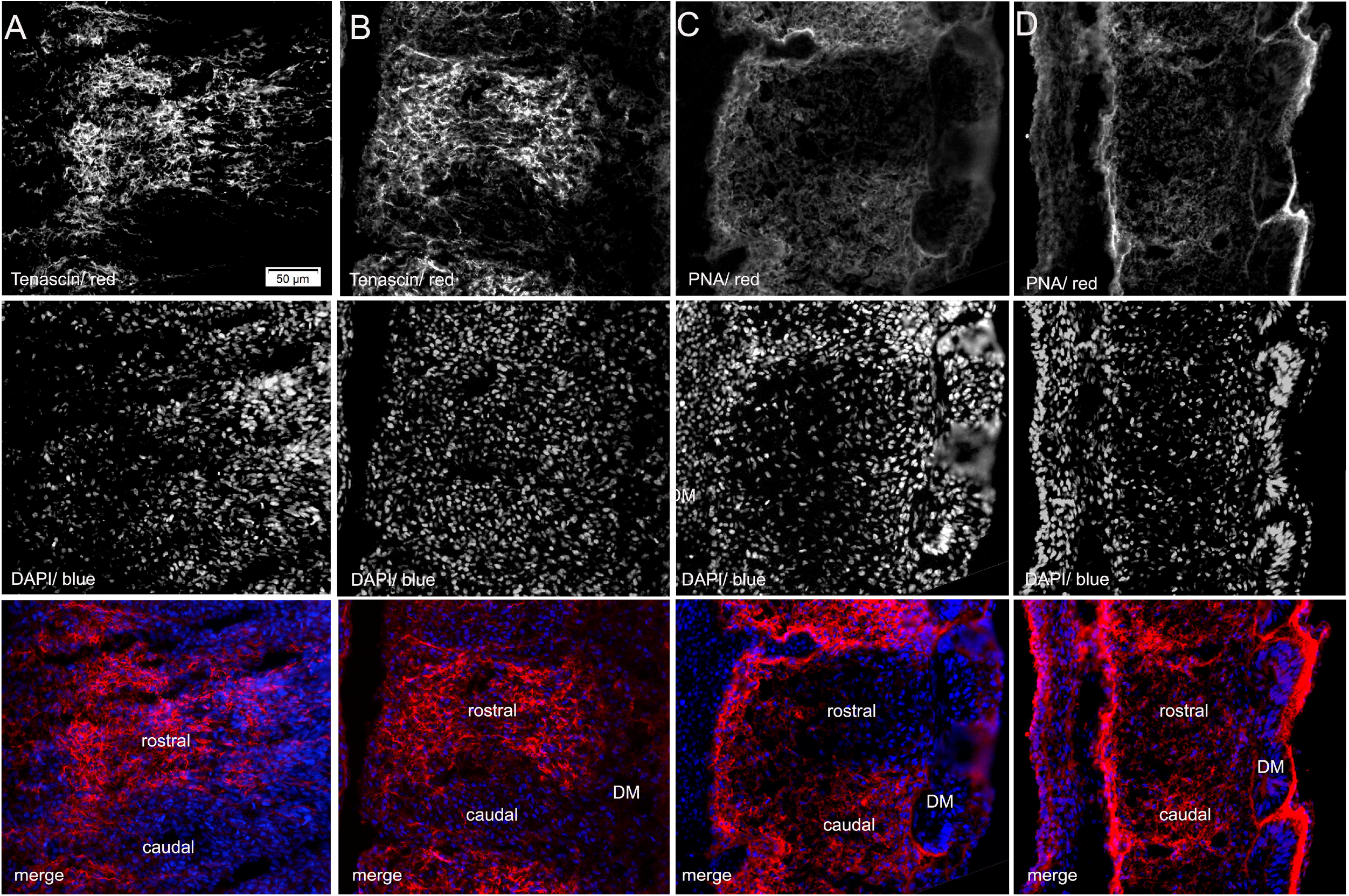
TGFβ signaling regulates rostral-caudal patterning in the sclerotome. (A, C) DMSO or (B,D) SB431542 injected E3.5 embryos were embedded in OCT, sectioned, and slides containing clear somite borders were selected. (A) DMSO control embryos (n=4) and (B) SB431542 (n=7) embryos were stained with a Tenascin antibody to label the rostral domain in somites. (C) DMSO control (n=3) embryos and (D) SB431542 embryos (n=4) were also stained with PNA to label the caudal domain in somites. (A’-D’). Hoechst was used as the counter stain. Left is ventral and right is dorsal. n= denotes the number of biological replicates. DM, dermomyotome.

## DISCUSSION

The importance of TGFβ signaling in the development of the spinal column has been shown in numerous studies (Baffi et al., 2004, Baffi et al., 2006, Cox et al., 2014, Ban et al., 2019). Phenotypes in mice with conditional deletion of *Tgfbr2* in sclerotome are consistent with defects in resegmentation although resegmentation was not tested directly (Baffi et al., 2004, Baffi et al., 2006). Here we directly tested the role of TGFβ signaling in sclerotome resegmentation during spinal column formation. We used lineage tracer dyes, DiD and DiO to follow somite derivatives including the sclerotome over time, and SB431542, a TGFβR1 inhibitor, to determine the role of TGFβ signaling in resegmentation and development of the spinal column. The chick model was chosen for this study due to its ability to develop and remain viable in *ex ovo* culture conditions. *Ex ovo* culturing of embryos permits easy manipulation and visualization of developmental processes (Aoyama and Asamoto, 2000, Christ et al., 2004, Huang et al., 2000). In addition, it has been demonstrated that the processes of resegmentation and sclerotome differentiation in chick are similar to that of mouse and human (Tanaka and Uhthoff, 1981, Takahashi et al., 2013). We established that SB431542 inhibited TGFβ signaling in chick somites at the concentrations used by looking at expression of known down-stream targets of TGFβ, most notably Scx (Clayton et al., 2020). We also showed a reduction in the AF of the IVD after treatment with SB431542, similar to what is seen in *Tgfbr2* conditionally deleted mice (Baffi et al., 2004) further supporting the use of chick as a model in these experiments.

Increased thickness of MF20 labeled myofibers was observed in SB431542 injected chick embryos. TGFβ signaling has been shown to act as an inhibitor of myofiber differentiation acting through Smad3 to inhibit MyoD, one of the master transcriptional regulators of muscle development (Kollias and McDermott, 2008, Liu et al., 2001, Filvaroff et al., 1994). TGFβ inhibited myoblasts were not able to fuse to form myotubes (Filvaroff 1994). In addition, treatment of limb bud organ cultures with neutralizing antibodies to TGF-β ligands resulted in the early appearance of large secondary myotubes, similar to what we observed here (Kollias and McDermott, 2008, Cusella-De Angelis et al., 1994). It was suggested that TGF-β prevents premature differentiation of migrating myoblasts to permit proper muscle formation (Kollias and McDermott, 2008). Our observations in chick further support this function of TGF-β in muscle development as well as supporting pharmacological inhibition of TGF-β signaling in the chick model.

Rostral-caudal polarity in the sclerotome has been shown to be important for IVD formation. “Rostralizing” or “caudalizing” mouse sclerotome through deletion of specific transcription factor markers of the rostral or caudal domains had severe effects on IVD differentiation and spinal column formation (Takahashi et al., 2013). For example, deletion of Mesp2, a marker of rostral sclerotome, or Ripply1/2, a marker of caudal sclerotome, resulted in misshapen vertebrae and missing IVDs supporting the importance of rostro-caudal identity in development of the spine (Takahashi et al., 2013). In the present study, we showed that resegmentation was completed by E6.5 in the chick and was disrupted in the presence of SB431542 as evidenced by the altered pattern of DiO and DiD labeled domains in the VBs. We noticed that the caudal labeled domains were shifted rostrally, with a smaller rostral domain in treated compared to controls. We hypothesized that this caudal shift was reflective of disrupted rostro-caudal polarity in earlier sclerotome. One of the most notable defects in Col2aCre; *Tgfbr2*^LoxP/LoxP^ mice was alterations in the polarity of the early sclerotome before resegmentation (Baffi et al., 2004 Baffi et al., 2006). Expansion of the caudal domain, as measured by Pax1/9 expression, anteriorly without alterations in localized expression of Tbx18, a rostral marker, in Col2aCre; *Tgfbr2*^LoxP/LoxP^ mice resulted in both rostral and caudal markers being co-expressed in the rostral half of the sclerotome (Baffi et al., 2006). Here, we used Tenascin as a marker of rostral sclerotome and PNA as a marker of caudal sclerotome. Treatment with SB431542 did not alter the expression domain of Tenascin; however, the expression domain of PNA was expanded through the entire sclerotome segment so that it overlapped with Tenascin. This unusual rostro-caudal pattern is similar to what we saw in Col2aCre;Tgfbr2^LoxP/LoxP^ mice (Baffi et al 2006). The occurrence of the same unique patterning in two different animal models supports the hypothesis that TGFβ regulates rostro-caudal polarity in the sclerotome.

It has been suggested that alterations in rostal-caudal polarity affect resegmentation (Takahashi et al., 2013). We propose that early alterations in rostro-caudal polarity in the sclerotome due to inhibition of TGFβ signaling contribute to the observed disruption in resegmentation. Alterations in rostro-caudal polarity and subsequent resegmentation would then be expected to alter the context in which cells differentiate, affecting development of the spinal column. In the chick, the IVD forms from the rostral domain of the sclerotome near von Ebner’s fissure, which is the boundary that separates the rostro-caudal domains (Bruggeman et al., 2012). Aberrant expression of caudal markers rostrally could be influencing how and where von Ebner’s fissure forms and thus cause alterations in resegmentation. In addition, co-expression of both rostral and caudal genes within the rostral domain in chick as a result of inhibition of TGFβ signaling could potentially cause alterations in differentiation pathways in the rostral domain leading to IVD malformations. Many domain markers are transcription factors initiating distinct differentiation protocols within their respective domains (Leitges et al., 2000, Morimoto et al., 2007, Kawamura et al., 2008). Inappropriate expression of these markers rostrally due to inhibition of TGFβ signaling could result in alterations in cell fate decisions that would affect development of the spinal column, and in this case the AF. Alterations in rostro-caudal polarity and subsequent resegmentation would also be expected to alter the context in which cells differentiate, affecting development of the spinal column. Alterations to cell fate decisions after resegmentation could then occur because cells are not in the correct locations to receive instructive, permissive, or competence signals.

In summary, this study provides the first direct evidence that TGFβ regulates resegmentation. We propose that TGFβ regulates rostro-caudal polarity in the sclerotome, which subsequently affects resegmentation and placement or differentiation of tissues within the spinal column.

## MATERIALS AND METHODS

### *Ex ovo* chick embryo culture

Specific Pathogen Free Premium Fertilized eggs (Charles River) were incubated at 99 degrees Fahrenheit and 55% humidity in a GQF 1500 series incubation cabinet (GQF Manufacturing Company) for 60-65 hours, 2 and a half days (E2.5), until embryonic stage HH16. After incubation, eggs were cracked and cultured in Deep Dish Petri Dishes (Fisherbrand) using previously published conditions (Auerbach et al., 1974, Yalcin et al., 2010) before being injected. Injected embryos were then returned to the incubator and allowed to grow until the indicated embryonic stage/ developmental day. Preestablished exclusion criteria included any embryo that had a birth defect on the day of injection and contamination or death of the embryo before the end point of the experiment. Statistics including T-tests and ANOVA were preformed to quantify various descriptive endpoints. Tests were run to assure that the data were normal and that variance was equal between each group using GraphPad Prism before the appropriate statistical test was selected. All graphed data is shown as individual data points.

### Embryo injections

All injections were performed using a Pli-100A pico-liter microinjector (Warner Instruments) at embryonic day 2.5 (E2.5). Selection for animals that were injected with either DMSO or SB431542 were randomized. The embryos analyzed on different developmental days for the microscopy experiments were matched for similar Hamburger and Hamilton (HH) stages to make a more accurate comparison between DMSO and SB431542 treatments. For skeletal preparations, western blot, and whole mount *in situ* analysis, eggs were cracked and injected with a control solution of DMSO or 50μM SB431542 Catalog #S1067 (Tocris) in thoracic somites 19 through 26. Somites were injected with solution until there was visible swelling in the somite. After injections, embryos were placed back into the incubator and cultured for 24 hours to reach embryonic day 3.5 (E3.5) for western blot, immunostaining, and whole mount *in situ* hybridization. Histology was done at E6.5 days. For skeletal preparations, embryos were cultured for 10 days post injections until embryonic day 12.5 (E12.5). For lineage tracing experiments, E2.5 embryos were injected with fluorescent, lipophilic dyes, DiD Catalog #D7757 and DiO Catalog #D257 (Thermofisher Scientific), with or without the presence of 50μM SB431542 into somites 21 through 26 on the right side only. DiD was injected into somites 21, 23 and 25, and DiO was injected into somites 22, 24 and 26. After injections, embryos were collected immediately after injections at E2.5 and then for 24hrs up to 4 days post injections until E6.5 for confocal imaging or histology experiments.

### Skeletal preparations

Injected embryos were allowed to reach E12.5 and then sacrificed. Embryos were removed from their extraembryonic membranes, decapitated, washed in Dubeccos phosphate buffered saline (DPBS) (Gibco), fixed for 1 hour in 4% paraformaldehyde (PFA), and then submerged in 70% ethanol overnight. The skin and organs were then removed before placing the embryos in the alcian blue solution (10mg X-Gal powder (Thermofisher Scientific), 20mL acetic acid and 80mL 95% ethanol) overnight (Nagy et al., 2009). Embryos were then rehydrated in 50% and then 25% ethanol: 0.5% potassium hydroxide washed before being stained with alizarin red (0.002% powder in 0.5% potassium hydroxide) for 24hours. Tissues were cleared in increasing concentrations of glycerol: 0.5% potassium hydroxide before being stored and imaged in 100% glycerol. Images were taken on an Olympus SZX12 microscope using a 0.5X PF objective. To measure IVD area changes, Image J was used to draw ROIs around the IVDs above thoracic vertebrae 5, 6, and 7 (T5, T6, T7) in each embryo, and area of each ROI was calculated. Differences between groups was analyzed using an unpaired two-tailed t-test in GraphPad Prism.

### Whole mount *in situ*

Injected embryos were allowed to reach E3.5, isolated, and washed in diethyl pyrocarbonate (DEPC) treated PBS. To perform whole mount in situ, the protocol reported in (Riddle et al., 1993) was used. The pBS cScx 3UTR plasmid Catalog #13957 (Addgene) created by (Schweitzer et al., 2001) was used as the template to create the Scx mRNA probe by utilizing the DIG RNA Labeling Kit Catalog #11175025910 (Roche) and the T3 RNA polymerase Catalog #11031163001 (Roche). Staining of the embryos was completed using BM-Purple substrate Catalog # 11442074001 (Roche), and imaging was done with an Olympus SZX12 microscope using a 0.5X PF objective.

### Western blot

Injected tissue was isolated from the thoracic region of embryos and lysed with Radio Immunoprecipitation Assay (RIPA) buffer (Roche). Total protein concentration was measured using a DC Protein Assay kit (Bio-Rad Laboratories) and 40μg of protein lysate per sample was loaded on 4-20% polyacrylamide gels (Bio-Rad Laboratories) for separation. Protein was then transferred to polyvinylidene fluoride membranes using a Trans-Blot Turbo Transfer system (Bio-Rad Laboratories). All membranes were blocked with 5% Bovine Serum Albumin (Sigma-Aldrich) and incubated with anti-SCXA antibody, Catalog #PA5-23943 (Invitrogen); anti-phospho Smad23 Catalog #8828S (Cell Signaling); anti-Smad3 Catalog #9513S (Cell Signaling); anti-Adamtsl2 Catalog #ab97603 (Abcam); and anti-α tubulin Catalog #200-301-880, (Rockland) overnight at 4 degrees Celsius. Membranes were washed with Tris-buffered saline containing 0.1% Tween 20 (TBST) and incubated with anti-Rabbit-HRP, Catalog #7074S (Santa Cruz Biotechnology) for 1 hour at room temperature. The chemiluminescence was detected by the Supersignal West Dura kit (Thermo Scientific). All images were acquired on a ChemiDoc MP system (Bio-Rad Laboratories), and quantification of blots was performed using ImageJ. Statistical analyses were performed using an unpaired, two-tailed t-test on GraphPad Prism. Asterisks denote p < 0.05

### Confocal microscopy and live-cell imaging

After indicated post injection incubation times, embryos were isolated and dissected. For E2.5 embryos the entire embryo was imaged, for E3.5 embryos the head was removed, for E4.5 and E5.5 embryos the head and limb buds were removed, and for E6.5 embryos the head, limb buds, and skin were removed to better visualize labeled cells. Embryos were placed in a glass bottom, cell imaging dish Catalog #0030740009 (Eppendorf) and submerged in imaging solution (80% egg white and 20% 1.2% UltraPure LMP Agarose Catalog #16500100 (Invitrogen) in Ringer’s solution (FisherScientific). Embryos were then taken to the microscopy core for 3D imaging on a Nikon Ti2 laser scanning confocal. For maximum intensity projections (maxIP), the 10x objective was used to capture images at a depth of 2.5-3.5 μm/z through the z plane for a total thickness of 150-200μm dependent upon the embryonic stage. Z stacks were combined to make a maxIP. For live-cell imaging, embryos were placed into a Tokai Hit incubation stage chamber at 37 degrees Celsius to maintain viability. Embryos were imaged every 10 minutes for 12 hours to produce 73 z stacks per embryo and the average z stack took 8-9 minutes. Z stacks were then combined into maxIP images and then looped together to make a video. To capture development of the spinal column from injections at E2.5 to resegmentation at E6.5, a total of 9 embryos underwent live-cell imaging, with a different embryo being imaged every 12 hours over the span of 4 days. All maxIP images and videos were made using the Nikon NIS-Elements Advanced Research software. Replicate numbers for each image and video are indicated in the figure legends. Videos are located in the supplemental figures section.

### Histology

The spinal column was dissected from isolated embryos, washed in PBS, and fixed for 1hr in 4% PFA. Embryos were placed in 30% sucrose overnight at 4 degrees Celsius and then placed in graded 30% sucrose: OCT solutions for 1 hour until reaching 100% OCT. Embryos were embedded in OCT and 10μm sections were made using a Lecia cryostat. Slides were fixed with 4% PFA for 20 minutes before immunofluorescence (IF) or histological staining. For alcian blue staining, slides were treated with alcian blue overnight in a humidified chamber at 4 degrees Celsius and then imaged with an Olympus SZX12 microscope using a 1X PF objective. For peanut agglutinin (PNA) staining, slides were permeabilized with 0.1%Trition X for 10 minutes and then blocked in 1% BSA for 30 minutes. PNA-Rhodamine (Vector Labs) was added to slides at a dilution of 1:100 for 1 hour. Immunofluorescence was conducted by permeabilizing with 0.1% TritonX and blocking in 1% Goat serum in TBST before adding the primary antibodies of anti-MF20 (DSHB) at a dilution of 7:250, anti-Tenascin (DSHB) at a dilution of 1ug/250ul. An anti-mouse Alexa 555 secondary was added at a dilution of 1:250. Indicated slides were treated with Hoechst 33258 (Invitrogen) at a dilution of 1:1000 before being mounted and imaged with an Olympus fluorescent microscope using a 20X PF objective. To measure myofiber fiber differences via MF20, ImageJ was used to zoom in and take images from the left, middle, and right of each image. The width of all myofibers, white brackets in panels B and D, in each zoomed in area was measured and average fiber thickness per sample was calculated.

## Acknowledgements

This study was funded by NIH and NIAMS R01 AR053860 to R.S and T32 AR069516 (PI Bridges) to SWC. A special thanks to Robert Grabski, PhD in the UAB High Resolution Imaging Facility, P30 AR048311, for the extensive training on the laser-scanning confocal that allowed the collection and analysis of the live-cell imaging data. Figures 1A, 3A, 4A, and 6A were created at Biorender.com.

## Author Contributions

S.W.C. and R.S. contributed to the conception and design of the study. S.W.C. and R.M. acquired the data. S.W.C. and R.S. contributed to the analysis and interpretation of the data, as well as writing the manuscript. All authors approved the final version of the manuscript and take responsibility for the integrity of the work.

