## Supplementary material for "TGFβ signaling is required for sclerotome resegmentation during development of the spinal column in *Gallus gallus*": Fig. S

### Supplemental Figures

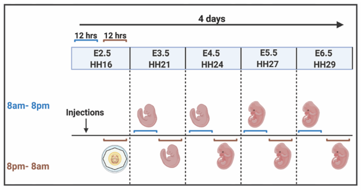


**Supplemental Fig. 1. Live imaging of labeled somites.** Embryos were injected with dyes at E2.5, isolated at indicated times, embedded in a yolk/agarose imaging solution to maintain viability as described in methods, and then imaged on a laser scanning confocal microscope in 12-hour (hr) intervals over the course of 4 days. A separate embryo was imaged every 12 hours. Changes in cell death as determined by Calcein-Ethidium cell staining were not observed (Fig S2). At E2.5 days, live cell imaging showed that the somites were well labeled (Movie 1) and that there was some movement of cells ventrally over the first 12 hrs after labeling (Movie 1). Dermamyotome which forms dorsal and lateral in the embryo was followed over time. In labeled DMSO treated embryos, the most striking observation was the appearance of elongated cells in the dorsal lateral aspect of the embryo starting at about E4.5 days (Movie 2). We hypothesized that these cells were developing myocytes based on their morphology which was confirmed by MF20 staining (Fig 3). SB431542 injected embryos are shown in Movie 3.

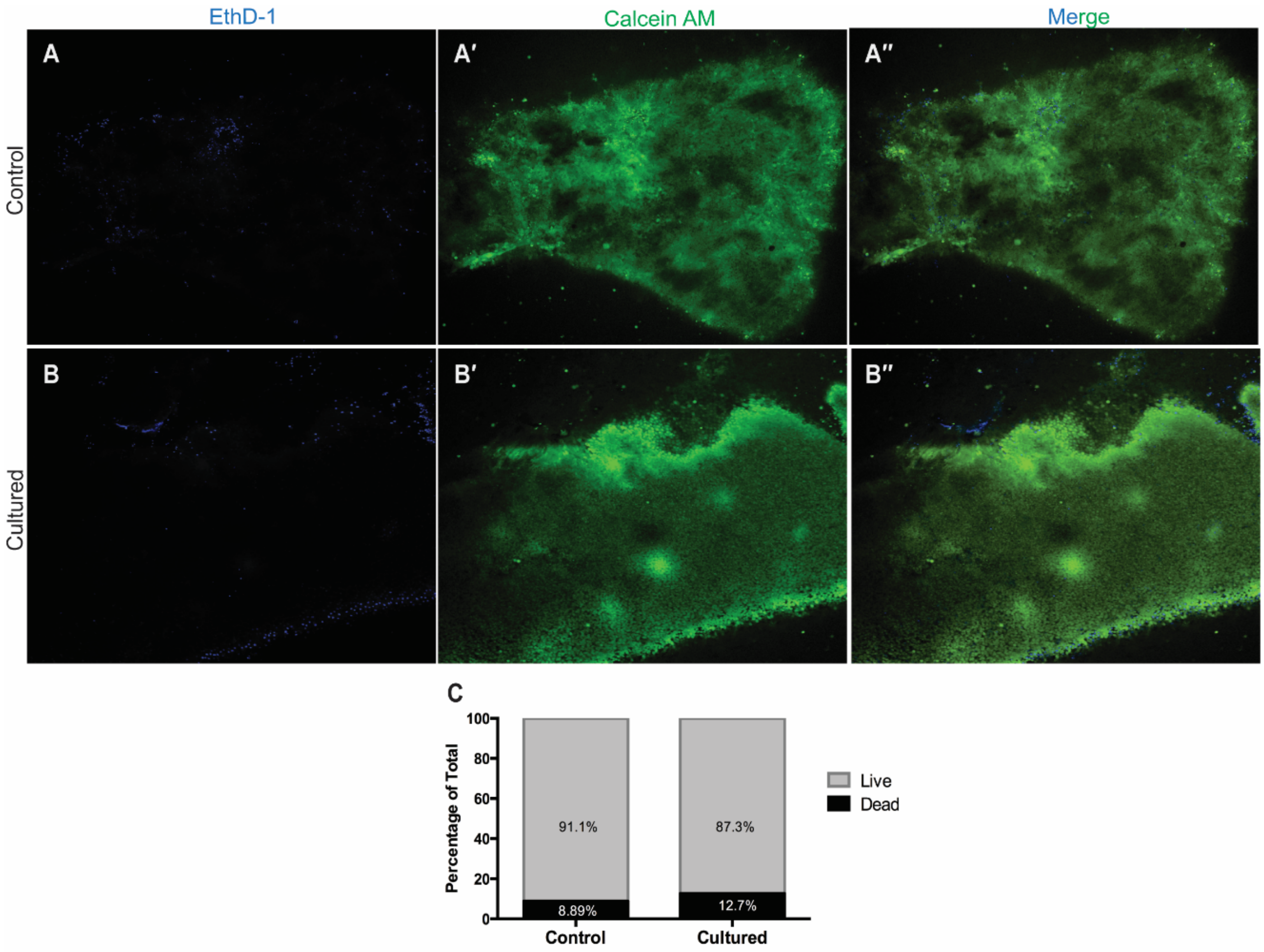


**Supplemental Fig. 2**: **Chick embryos are still viable after 12hrs cultured in artificial conditions.** (A) E4.5 control embryos (n=3) that were allowed to remain within *ex ovo* conditions or (B) placed in a 1:1 yolk: agarose imaging solution and imaged (n=3) for 12hrs and then stained with a Live/Dead kit. EthD-1 stains dead cells blue and Calcein AM stains living cells green. Images were taken at a superficial, medial, and deep depth of each embryo and the Image J particles tool was used to count labeled cells from each channel at each depth. The average number of live/dead cells was calculated per embryos and (C) an ANOVA was ran to between controls and cultured embryo values. A Tukey's post hoc analysis was ran to compare differences between groups. The control and cultured embryos had similar percentages of live to dead cells and there was no statistically significant increase in cell death in cultured embryos. And, in both conditions there was a significantly larger portion of living cells rather than dead cells.

**Movie 1: 12hr live cell imaging of an E2.5 control embryo.** Imaged from 8pm-8am. N=1. Left is ventral. Right is dorsal. Top is anterior. Bottom is posterior. 10x

**Movie 2: 12hr live cell imaging of an E4.5 control, DMSO injected embryo.** Imaged from 8pm-8am. N=2. Left is ventral. Right is dorsal. Top is anterior. Bottom is posterior. 10x

**Movie 3: Myofiber formation in an E4.5 SB431542 injected embryo during live cell imaging.** Imaged from 8pm-8am. N =1. Left is ventral. Right is dorsal. Top is anterior. Bottom is posterior. 10x

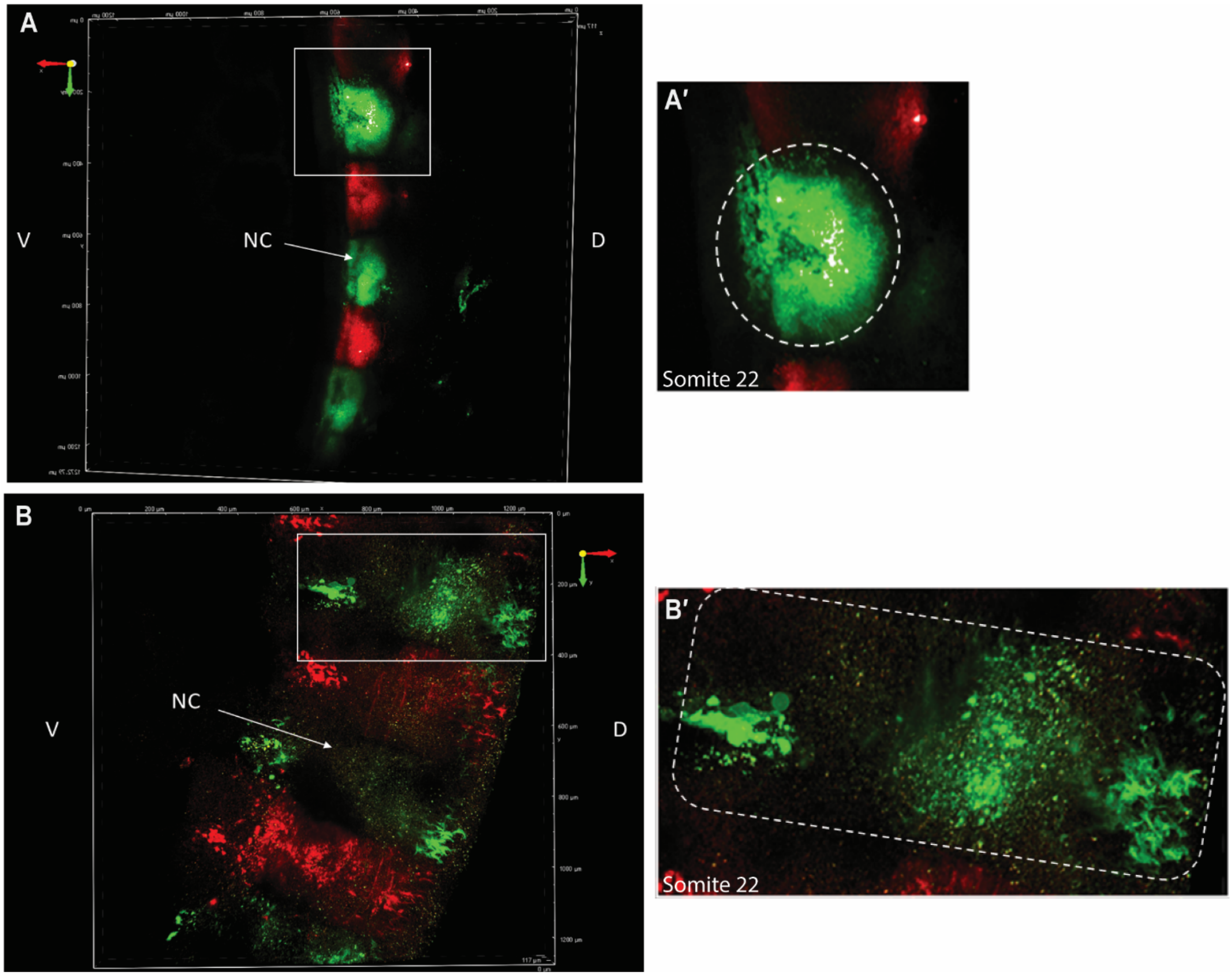


**Supplemental Fig. 3: Somite undergo an epithelial to mesenchymal transition (EMT) between E2.5 and E3.5.** (A) DiD and DiO labeled somites are intact and spherical in shape at E2.5, indicating they have not yet begun to disperse due to EMT. (A′) A zoomed in image of somite 22, outlined in a white box in panel A, showed a closer look at the spherical shape of somites at E2.5. (B) At E3.5, somites no longer have a discernable spherical shape and labeled cells have begun to migrate. (B′) A zoomed in image of what was somite 22 showed how migration has occurred over 24hrs since somites were injected at E2.5. Distance in the X and Y planes are labeled on the X and Y axes in microns.

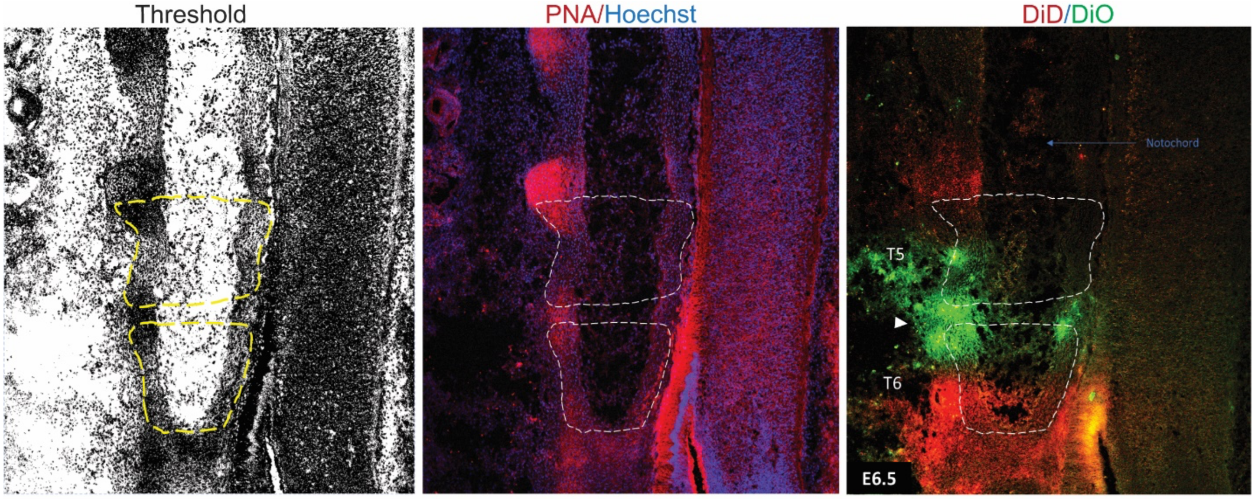


**Supplemental Fig. 4**: **Example of the thresholding technique used to outline vertebrae.** The Image J threshold tool was used to decrease background noise on PNA stained images and then the ROI function was used to outline VB area. VB outlines were drawn on the threshold images and then super imposed onto DiD/DiO stained images to determine VB area more accurately.
