## Supplementary figures and images for "TGFβ signaling is required for sclerotome resegmentation during development of the spinal column in *Gallus gallus*"

### Movie 2

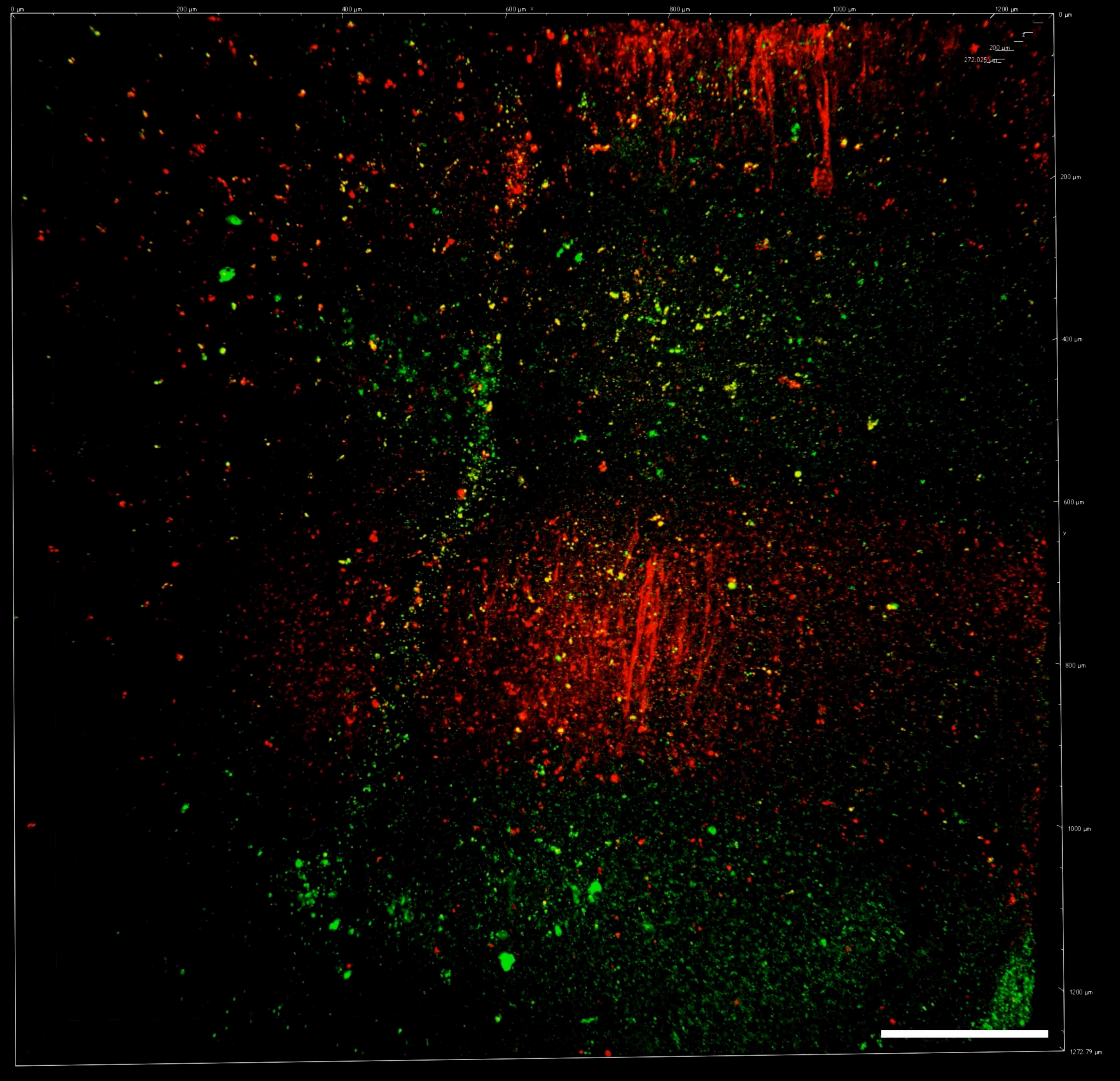

### Movie 3

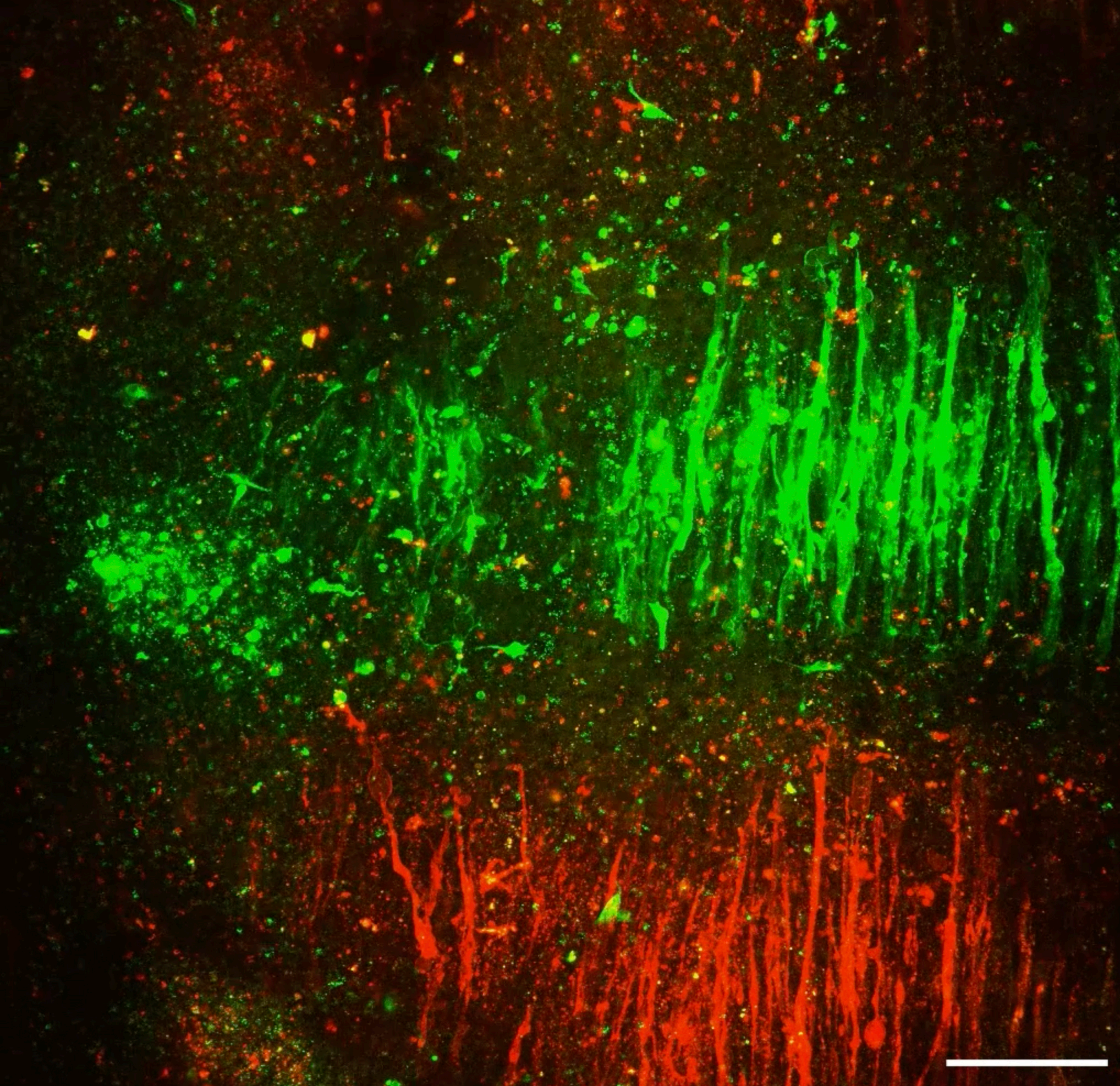

### Movies 1

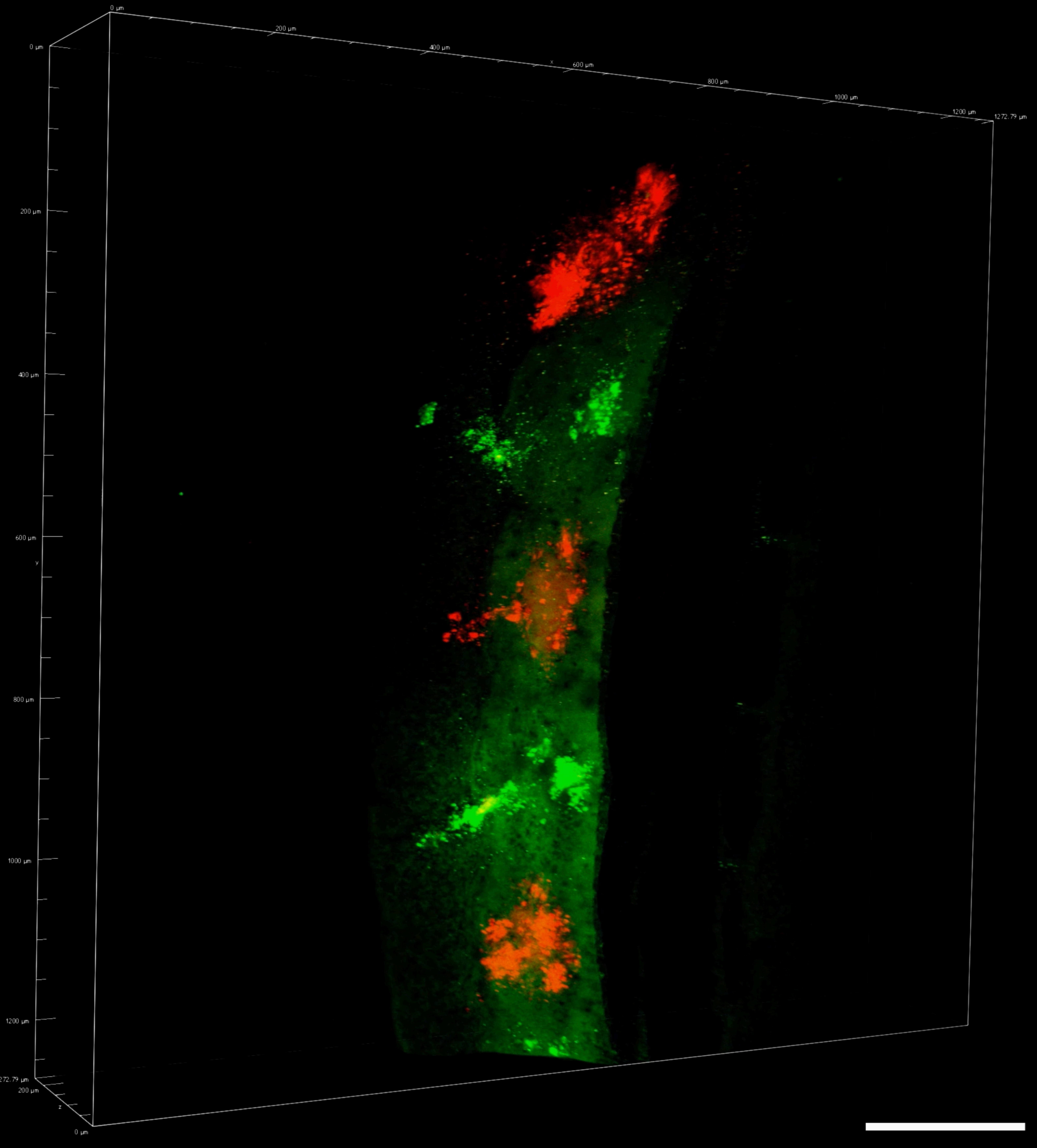
